# The RXLR-EER Motif Determines an Unconventional Secretion Pathway Associated with Extracellular Vesicle Production

**DOI:** 10.1101/2025.07.07.663508

**Authors:** Wei Wang, Shumei Wang, Lin Xu, Lydia Welsh, Petra C Boevink, Steve C Whisson, Paul RJ Birch

## Abstract

*Phytophthora infestans*, the cause of potato late blight disease, delivers a suite of RXLR effectors into host plant cells to subvert immunity, whereas apoplastic effectors act extracellularly. Although the RXLR-EER motif in these effectors is critical for host translocation and is cleaved prior to secretion, the relevance of this processing is poorly understood. Prior evidence suggests RXLR effectors utilize a distinct, unconventional secretion pathway, raising the question of whether the RXLR-EER motif influences selection of the secretion route. Here, we combined genetic, molecular and cell biology approaches to investigate the secretion pathway of RXLR effectors. Confocal microscopy revealed that RXLR and apoplastic effectors localize to distinct vesicular compartments in cultured hyphae. Moreover, fusing the ER retention signal KDEL to RXLR effectors did not impair their secretion, in contrast to apoplastic effectors, which were retained in the endomembrane system, indicating that RXLR effectors bypass the canonical ER-to-Golgi pathway. Importantly, RXLR effectors associate with extracellular vesicles (EVs), whereas RXLR-EER motif mutants show reduced EV association and are rerouted through the ER-to-Golgi secretion pathway. These findings demonstrate that the RXLR-EER motif governs effector sorting into an unconventional, EV-linked secretion route. This study sheds light on the molecular basis of effector trafficking in *P. infestans* and underscores the potential role of EVs in delivering virulence factors during host colonization.

## Introduction

The oomycete *Phytophthora infestans* is the causative agent of potato late blight, a devastating disease that triggered the Irish potato famine in the 19th century and continues to pose a major threat to global food security ^1, 2^. As a highly adapted hemi-biotrophic pathogen, *P. infestans* relies on a large and complex suite of effector proteins to manipulate host cell physiology and suppress immune responses, enabling successful colonization and disease progression.

The effectors can be broadly divided into two classes based on their site of action: apoplastic effectors, which are secreted into the extracellular space (apoplast) ^3^, where they act on the outside of plant cells, and cytoplasmic effectors with an Arg-any amino acid-Arg-Leu (RXLR) motif, downstream of a secretion signal peptide, often followed closely by EER (Glu-Glu-Arg) motif, which are encoded by hundreds of genes within the genomes of *Phytophthora* spp. and downy mildew pathogens ^4, 5, 6^. RXLR effectors are delivered into the host cell cytosol and target multiple proteins and processes at diverse subcellular locations to suppress immunity ^7, 8, 9, 10^. Whereas apoplastic effectors may be perceived by host pattern recognition receptors (PRRs) at the cell surface, RXLR effectors interfere with intracellular immunity by targeting various host processes within plant cells and can, in turn, be perceived by nucleotide-binding leucine-rich repeat receptors ^11^. The RXLR-EER motif is crucial for host cell translocation and is proteolytically cleaved prior to secretion, although the functional consequences of this processing are poorly understood ^12, 13, 14^.

Previous studies show that apoplastic effectors in *P. infestans* are secreted via the conventional ER-to-Golgi pathway, as indicated by their sensitivity to the inhibitor brefeldin A (BFA). In contrast, RXLR effectors are largely BFA-insensitive, suggesting they follow an unconventional secretion route^15, 16^. A similar dichotomy in effector secretion has been observed in the fungal pathogen of rice, *Magnaporthe oryzae*, where cytoplasmic effectors are delivered via a BFA-insensitive, non-classical pathway, whereas apoplastic effectors rely on BFA-sensitive ER-to-Golgi trafficking ^17^. However, it is important to recognise that BFA treatment can have off-target or pleiotropic effects, including disruption of endosomal trafficking, vesicle recycling, and broader alterations in membrane dynamics ^18^. These side-effects may complicate interpretation of BFA sensitivity assays, particularly in complex filamentous pathogens such as *P. infestans*, where multiple secretory routes may coexist. Therefore, while BFA sensitivity provides a useful initial indication of secretory pathway involvement, additional genetic and imaging-based approaches are essential to unambiguously distinguish between conventional and unconventional secretion routes.

Recently, emerging evidence suggests that RXLR effectors are secreted via an unconventional pathway potentially associated with production of extracellular vesicles (EVs)^19^. EVs are membrane-bound nanoparticles released by cells into the extracellular milieu and have been increasingly recognized as vehicles for intercellular communication and molecular cargo delivery between organisms from different kingdoms^20^. Recently, EVs have been shown to play a pivotal bidirectional transport role in plant–pathogen interactions ^21, 22, 23, 24, 25^. In addition to the intriguing link between EVs and effector trafficking, the mechanisms by which RXLR effectors are sorted to EVs, and the signals that determine their route of secretion, are largely unknown. In particular, whether the RXLR-EER motif influences intracellular sorting within the pathogen secretory systems has yet to be resolved.

In this study, we combined molecular, genetic and live-cell imaging approaches to investigate the secretion pathway of RXLR effectors in *P. infestans*. To determine whether these effectors utilize the classical ER–to-Golgi secretion route, we fused a canonical ER retention signal, KDEL (Lys-Asp-Glu-Leu), to RXLR effectors. The KDEL motif, typically found at the C-terminus of ER-resident proteins, is recognized by KDEL receptors in the cis-Golgi, which mediate the retrograde transport of these proteins back to the ER. While apoplastic effectors were largely retained in the ER after the addition of KDEL, RXLR effectors remained efficiently secreted, suggesting that they bypass the conventional ER–to-Golgi pathway. Furthermore, site-directed mutagenesis of the RXLR-EER motifs redirected these effectors to the classical ER-to-Golgi secretion route and disrupted their association with EVs, indicating that the RXLR-EER motif plays a role in effector trafficking within the pathogen.

Our findings provide compelling evidence that the RXLR-EER motif governs effector trafficking within *P. infestans*, channelling them into an unconventional secretory route linked to EV production. This study deepens our understanding of effector biogenesis and secretion and highlights EVs as potential mediators of virulence protein delivery in plant-pathogen interactions.

## Results

### RXLR effectors and apoplastic effectors are associated with different types of trafficking vesicles

Previous studies have shown that the secretion of RXLR effectors in *P. infestans* is largely insensitive to BFA, whereas the secretion of apoplastic effectors is BFA-sensitive. This differential response suggests that RXLR and apoplastic effectors utilize distinct secretion pathways ^15, 16^. To further explore this hypothesis, we co-expressed two apoplastic effectors, PiSCR74 and PiEPIC1, each C-terminally tagged with fluorescent proteins (mCitrine or mCherry respectively), in *P. infestans* and analyzed their localization using confocal microscopy. The two proteins exhibited substantial co-localization in puncta in cultured hyphae, consistent with the previous observations^15, 16^ that they are both trafficked through the conventional ER-to-Golgi secretory pathway (Fig. 1a). We have shown that RXLR effectors Pi04314 and Pi09216 strongly co-localized at vesicle-like structures in cultured hyphae^19^. Consistent with these findings, co-expression of Pi04314 and PiAVRblb1 also revealed robust co-localization at puncta in hyphae, reinforcing the concept that RXLR effectors follow a common intracellular trafficking route (Supplementary Fig. 1). In contrast, when apoplastic effector PiSCR74 was co-expressed with RXLR effector Pi04314, the two proteins localized to distinct vesicle-like compartments, providing supporting evidence that RXLR and apoplastic effectors are sorted through different secretory routes (Fig. 1b).

**Fig. 1.**
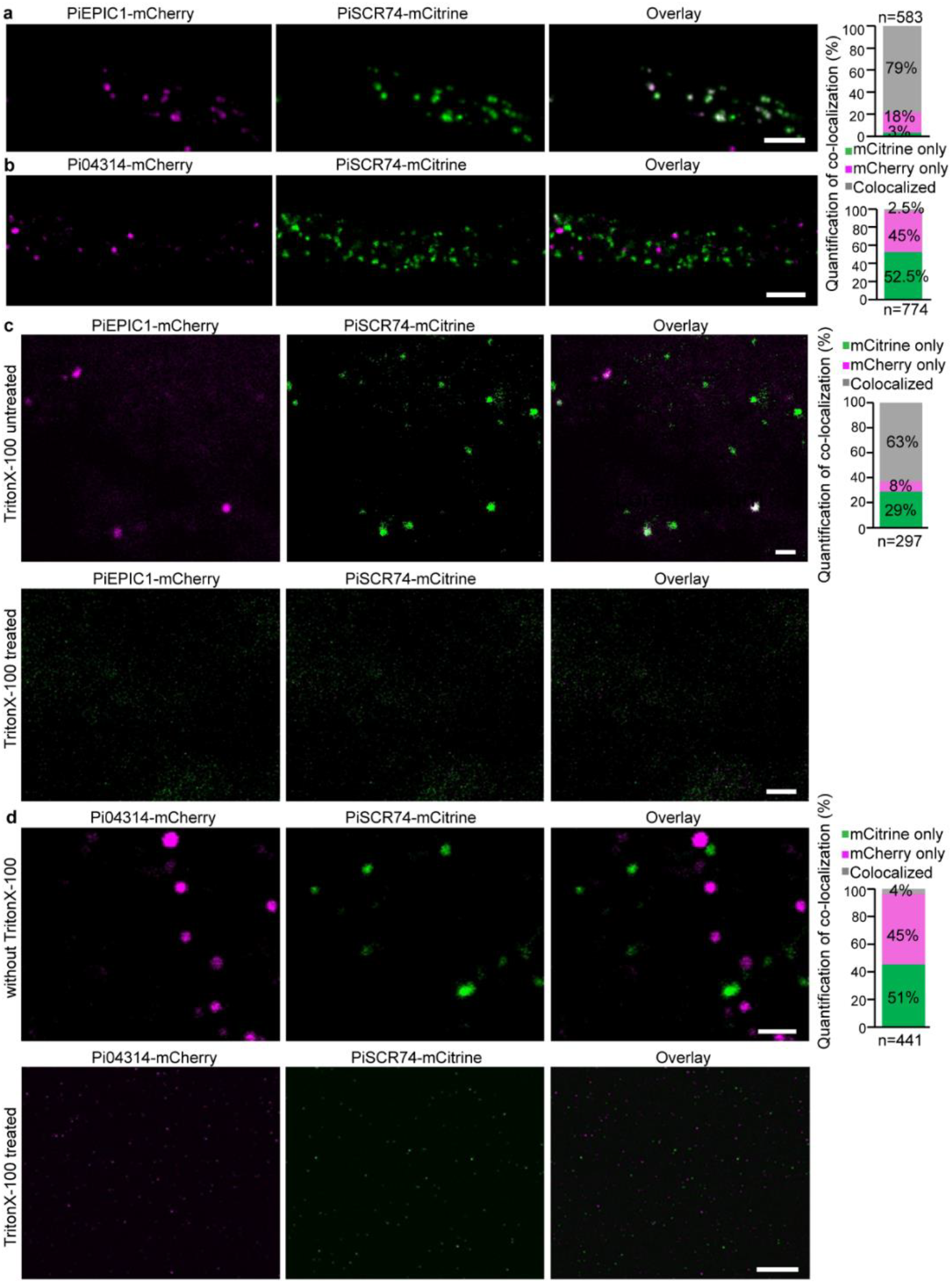
RXLR effectors and apoplastic effectors are associated with different types of secretory vesicles. **a**, Confocal microscopy images showing colocalization of PiSCR74-mCitrine with PiEPIC1-mCherry in puncta in cultured hyphae. **b**. PiSCR74-mCitrine co-expressed with the RXLR effector Pi04314-mCherry, showing a lack of colocalization in puncta in cultured hyphae. Scale bars, 10 µm (**a, b**). **c**, Endosomal vesicles isolated from the same dual transformants shown in **a** confirm colocalization between the apoplastic effectors (upper panel). Treatment with 1% Triton X-100 disrupts vesicle integrity, indicating that the punctate structures are membrane-bound (lower panel). Quantification of colocalization is shown in the accompanying graphs. **d**, Endosomal vesicles isolated from the same dual transformants as in **b** show almost no colocalization between Pi04314 and PiSCR74. Treatment with 1% Triton X-100 confirms both effectors are associated with membrane structures. Scale bars, 2 µm (**c**–**d**). Quantification of colocalization is presented in the accompanying graphs. The numbers of total and co-labelled puncta were determined using the DiAna plugin in Fiji (**a**–**d**). n=total puncta.

To investigate whether these effectors are associated with distinct populations of endosomal vesicles, we isolated endo-vesicles from cultured mycelia using a sucrose gradient that was bottom-loaded to allow vesicles to rise to their buoyant density and thus exclude protein aggregates (Supplementary Fig. 2a). Transmission electron microscopy of the vesicle fractions (VFs) confirmed the presence of typical membrane-bound vesicles (Supplementary Fig. 2b). Confocal imaging of the VFs revealed that fluorescent protein tagged PiSCR74 and PiEPIC1 largely co-localized within the same vesicle population. Furthermore, fluorescence signals from both effector fusions were abolished by pre-treatment with 1% TritonX-100 detergent, which disrupts lipid membranes but not protein aggregates^26, 27^ (Fig. 1c). Similarly, two fluorescently tagged RXLR effectors Pi04314 and PiAVRblb1 largely co-localized within the same vesicle population (Supplementary Fig. 2c). In contrast, Pi04314 and PiSCR74 fusions localized to separate vesicle populations. Similarly, their respective fluorescence signals were also detergent-sensitive, indicating membrane association (Fig. 1d). Immunoblot analysis confirmed that apoplastic and RXLR effectors were all present as intact fusion proteins in the vesicle fractions, and that detergent treatment abolished their presence in VFs (Supplementary Fig. 2d). These results indicate their association with membrane-bound vesicular structures, supporting the conclusion that RXLR and apoplastic effectors are sorted into distinct vesicle types and follow different secretion pathways in *P. infestans*.

**Fig. 2.**
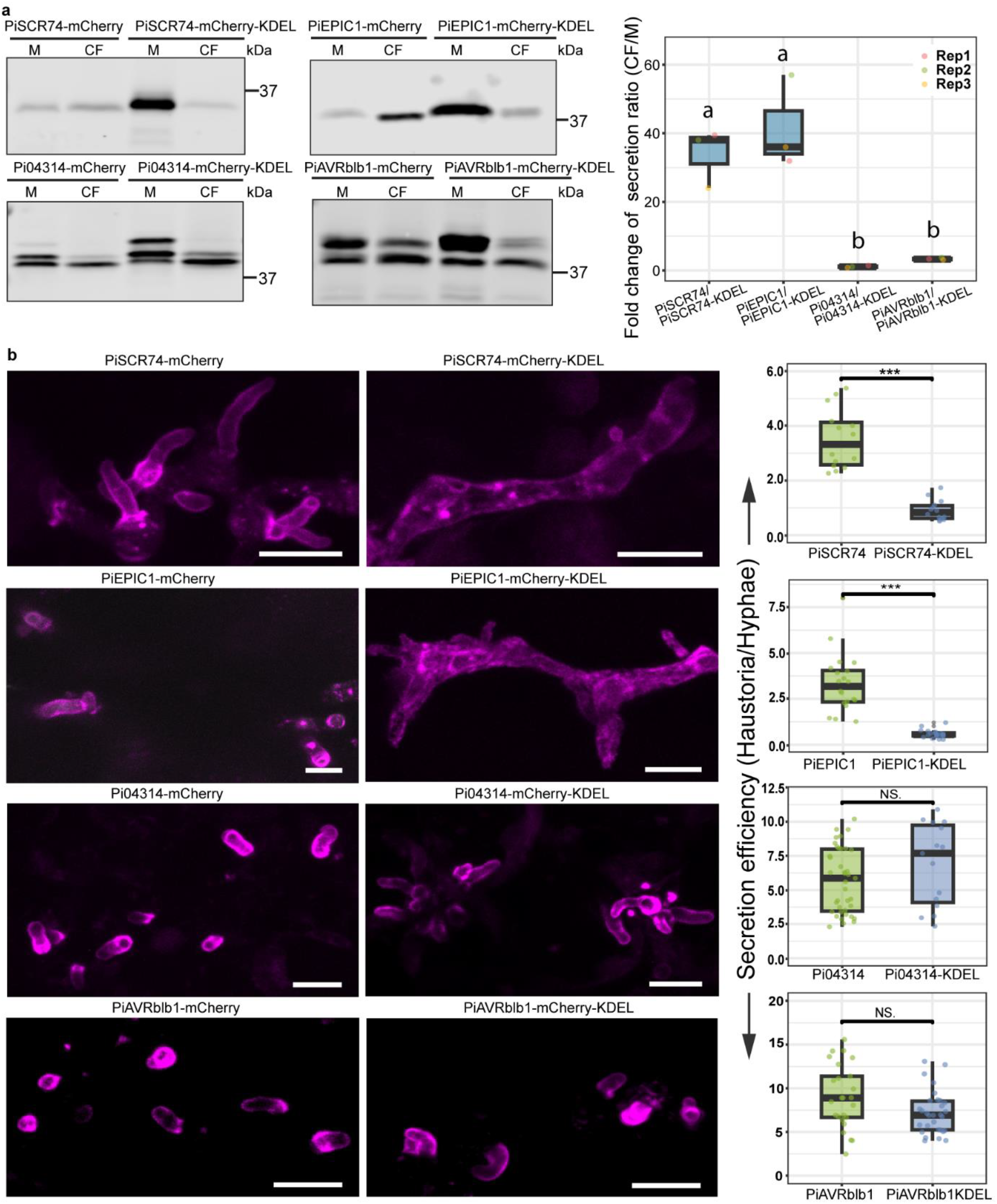
Apoplastic effectors secreted via Golgi but RXLR effectors take an alternative route. **a**, Immunoblot analysis shows retention of apoplastic effector fusions PiSCR74-mCherry and PiEPIC1-mCherry inside mycelia when fused to a KDEL motif, compared to efficient secretion in their unmodified counterparts. In contrast, the secretion efficicy of RXLR effector fusions Pi04314-mCherry and PiAVRblb1-mCherry was not significantly affected by the addition of the KDEL motif. Protein sizes are indicated in kDa. Quantification of the results is presented in the accompanying boxplot, highlighting the significantly greater impact of the KDEL motif on the secretion of apoplastic effectors compared to RXLR effectors. Secretion efficiency was calculated as the ratio of protein intensity in culture filtrate (CF) to that in mycelial extracts (M). Fold changes represent the difference in CF/M ratios between unmodified and KDEL-containing versions. Data are presented as mean ± SD from three biological replicates indicated by small solid circles. Statistical analysis was performed using one-way ANOVA with post-hoc Tukey HSD test to determine significance. **b**, Confocal microscopy images of infectious hyphae in potato leaf tissue showing that the accumulation of PiSCR74-mCherry and PiEPIC1-mCherry at haustoria is markedly reduced when appended with a KDEL motif, compared to their unmodified versions *in planta*. In contrast, accumulation of Pi04314-mCherry and PiAVRblb1-mCherry at haustoria is not significantly affected by the addition of KDEL. Quantification of the results is presented in the accompanying box plots. Secretion efficiency was calculated based on the fluorescence intensity ratio between haustoria and adjacent hyphae area. Data are presented as mean ± SD from three biological replicates. Individual data points are represented by small solid circles. Statistical analysis was performed using student’s *t*-test. Scale bars: 10 µm.

### RXLR effectors follow a Golgi-independent route after cleavage

Given that apoplastic effectors are secreted via the conventional ER-to-Golgi pathway and that RXLR effectors are insensitive to BFA treatment, we hypothesized that RXLR effectors utilize a Golgi-bypass secretion mechanism. To test this hypothesis, we employed a molecular strategy involving the addition of an ER retention signal—KDEL (Lys-Asp-Glu-Leu)—to the C-terminus of effector-fluorescent protein fusions. The KDEL motif is recognized by KDEL receptors (KDELRs) located in the cis-Golgi, which mediate retrograde transport of KDEL-tagged proteins back to the ER ^28^. Proteins secreted via a Golgi-independent route would be expected to escape this retention mechanism. To evaluate this, apoplastic (PiSCR74 and PiEPIC1) or RXLR (Pi04314 and PiAVRblb1) effector fluorescence protein fusions either with or without the KDEL motif, were expressed in *P. infestans* strains cultured in liquid medium. Immunoblotting revealed that apoplastic effectors were efficiently secreted under normal conditions, but their secretion was markedly reduced upon addition of KDEL, whereas the secretion of RXLR effectors was largely unaffected, indicating that their export does not primarily depend on Golgi-mediated trafficking (Fig. 2a).

We validated these findings *in planta* using confocal microscopy. All effectors accumulated at the haustorial interface under normal conditions. However, upon KDEL addition, apoplastic effectors accumulated in ER-like endomembrane compartments, consistent with receptor-mediated retention. In contrast, RXLR effectors accumulated at the haustorial interface even when fused to KDEL (Fig. 2b). These results strongly support the conclusion that apoplastic effectors are secreted through a conventional ER-Golgi pathway, whereas RXLR effectors are exported via a Golgi-bypass mechanism.

Our previous work showed that the RXLR-EER motif is not recognized or correctly cleaved in plant cells^14^. To determine whether proteolytic processing of this motif influences the pathway of secretion, the RXLR effector Pi04314 — with or without its native signal peptide (SP) — was transiently expressed in *Nicotiana benthamiana*. Confocal microscopy revealed that the SP-containing version was secreted into the apoplast, whereas the version lacking the SP localized to the nucleus or nucleolus, the expected site of the effector activity *in planta* ^29, 30^. While BFA treatment disrupted secretion of SP-tagged Pi04314, causing it to accumulation in BFA bodies^31^, it did not alter the subcellular localization of NSP-Pi04314 (Supplementary Fig. 3a). Addition of a C-terminal KDEL motif caused SP-04314-RFP to be retained in the ER (Supplementary Fig. 3b), confirming that uncleaved RXLR effectors follow the conventional ER-to-Golgi pathway in plant cells. These results highlight the importance of appropriate perception and cleavage of the RXLR-EER motif in directing these effectors toward an alternative, Golgi-bypass secretion route.

**Fig. 3.**
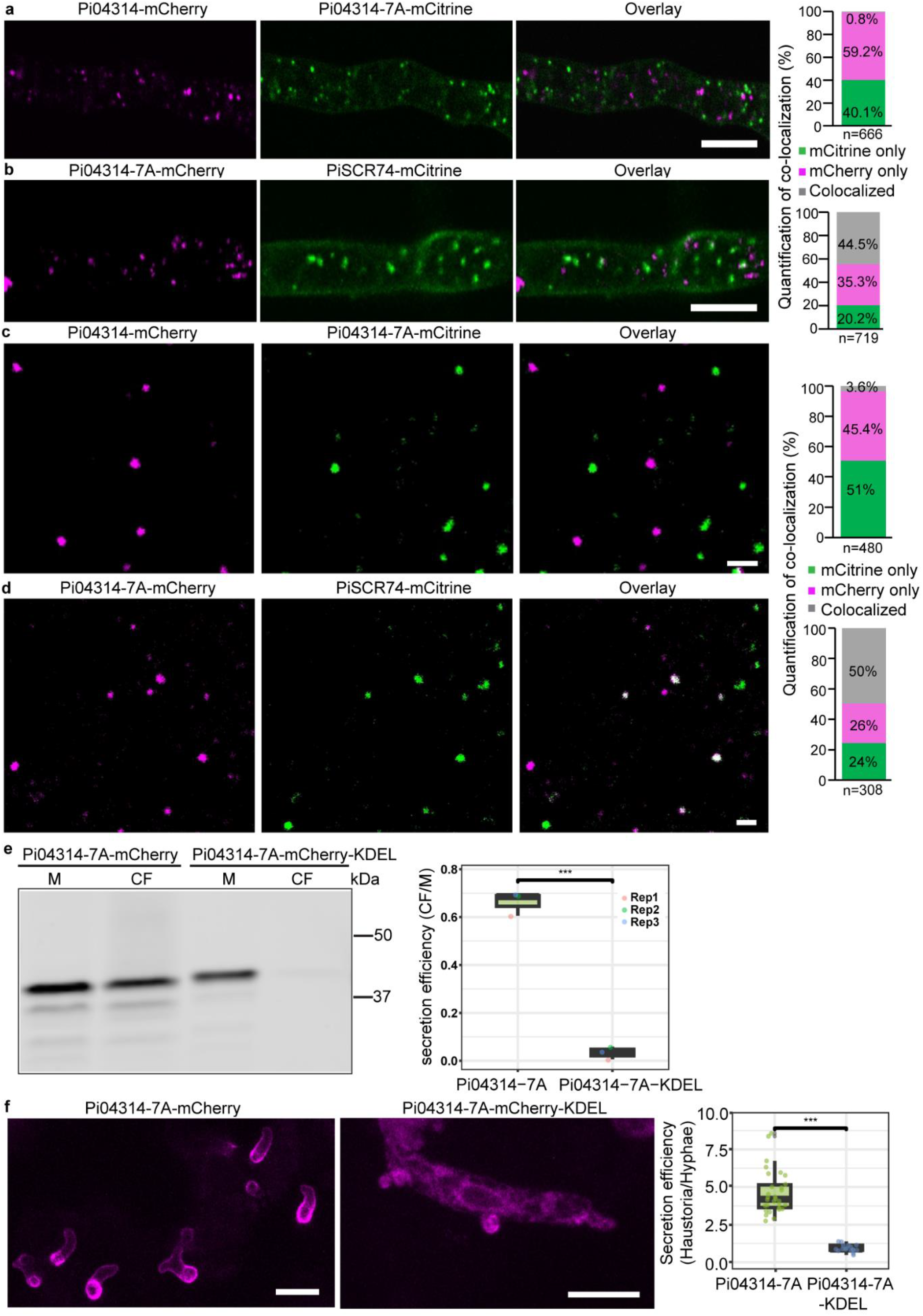
RXLR-EER motifs determine secretion pathways. **a**, Confocal microscopy images showing wild-type Pi04314-mCherry does not colocalize with the RXLR mutant effector Pi04314-7A-mCitrine. **b**, In contrast, colocalization of PiSCR74-mCitrine with Pi04314-7A-mCherry was observed in cultured hyphae. Scale bars, 10 µm. **c**, Endosomal vesicles isolated from the same dual transformants as in **a** confirm that Pi04314-7A does not colocalize with wild-type Pi04314. **d**, Confocal microscopy shows colocalization of Pi04314-7A with the apoplastic effector PiSCR74. Quantification of colocalization is shown in the accompanying graphs. The numbers of total (n) and co-labelled puncta were determined using the DiAna plugin in Fiji (**a**-**d**). Scale bars, 2 µm. **e**, Immunoblot analysis reveals that Pi04314-7A-mCherry-KDEL is largely retained in the mycelia (M) while secretion of the unmodified variant is readily detected in the culture filtrate (CF). Secretion efficiency was determined by calculating the ratio of protein abundance in the culture filtrate (CF) relative to that in M. Quantification of the protein secretion efficiency is presented in the accompanying boxplot. Data are presented as mean ± SD from three biological replicates indicated by small solid circles. Protein sizes are indicated in kDa. **f**, Confocal microscopy images of infectious hyphae in potato leaf tissue show that Pi04314-7A-mCherry secretion at haustoria is significantly diminished when fused to a KDEL motif, compared to the intense haustorial signal observed with the unmodified version *in planta*. Quantitative analysis is shown in the accompanying boxplots, where secretion efficiency was assessed by calculating the fluorescence intensity ratio between haustoria and hyphae. Data are expressed as mean ± SD from three independent biological replicates, with individual measurements represented by solid dots. Statistical significance was determined using student’s *t*-test. *** indicates p<0.001 (**e, f**). Scale bars, 10 µm.

### The RXLR-EER motif determines the secretion pathway

Given that the RXLR-EER motif functions as a protease processing site prior to secretion ^13, 14^, we sought to investigate whether this motif also determines the secretion route of RXLR effectors. In previous work, we showed that RXLR-EER motif mutants could still be secreted at haustoria^14^, but it is unclear if the secretion pathway was altered. To test this, we generated a mutant version of the RXLR effector Pi04314 in which the RXLR-EER motif was substituted with alanines (Pi04314-7A). Both wild-type (WT) and mutant versions were fused with different fluorescent proteins (mCherry or mCitrine) and co-expressed in *P. infestans*. Confocal microscopy of cultured hyphae revealed that the WT and mutant effectors localized to distinct vesicular compartments (Fig. 3a). Notably, the Pi04314-7A mutant co-localized significantly with apoplastic effector PiSCR74 (Fig. 3b), suggesting a shift in its trafficking route. Endo-vesicle isolation from the same transformants confirmed that the RXLR-EER mutant was sorted into vesicles distinct from those carrying the wild-type RXLR effectors (Fig. 3c), but largely co-localized with vesicles associated with apoplastic effector PiSCR74 (Fig. 3d). Importantly, addition of a KDEL motif to the RXLR-EER 7A mutants, for both Pi04314 and PiAVRblb1, significantly impaired their secretion into the culture medium (culture filtrate, CF) (Fig. 3e, Supplementary Fig. 4a). *In planta*, these KDEL-tagged mutants displayed ER retention patterns similar to those observed for KDEL-tagged apoplastic effectors (Fig. 3f, Supplementary Fig. 4b). These findings indicate that the RXLR-EER motif plays a key role in directing RXLR effectors into an unconventional, Golgi-independent secretion pathway, and its mutation leads to conventional ER-to-Golgi secretion.

**Fig. 4.**
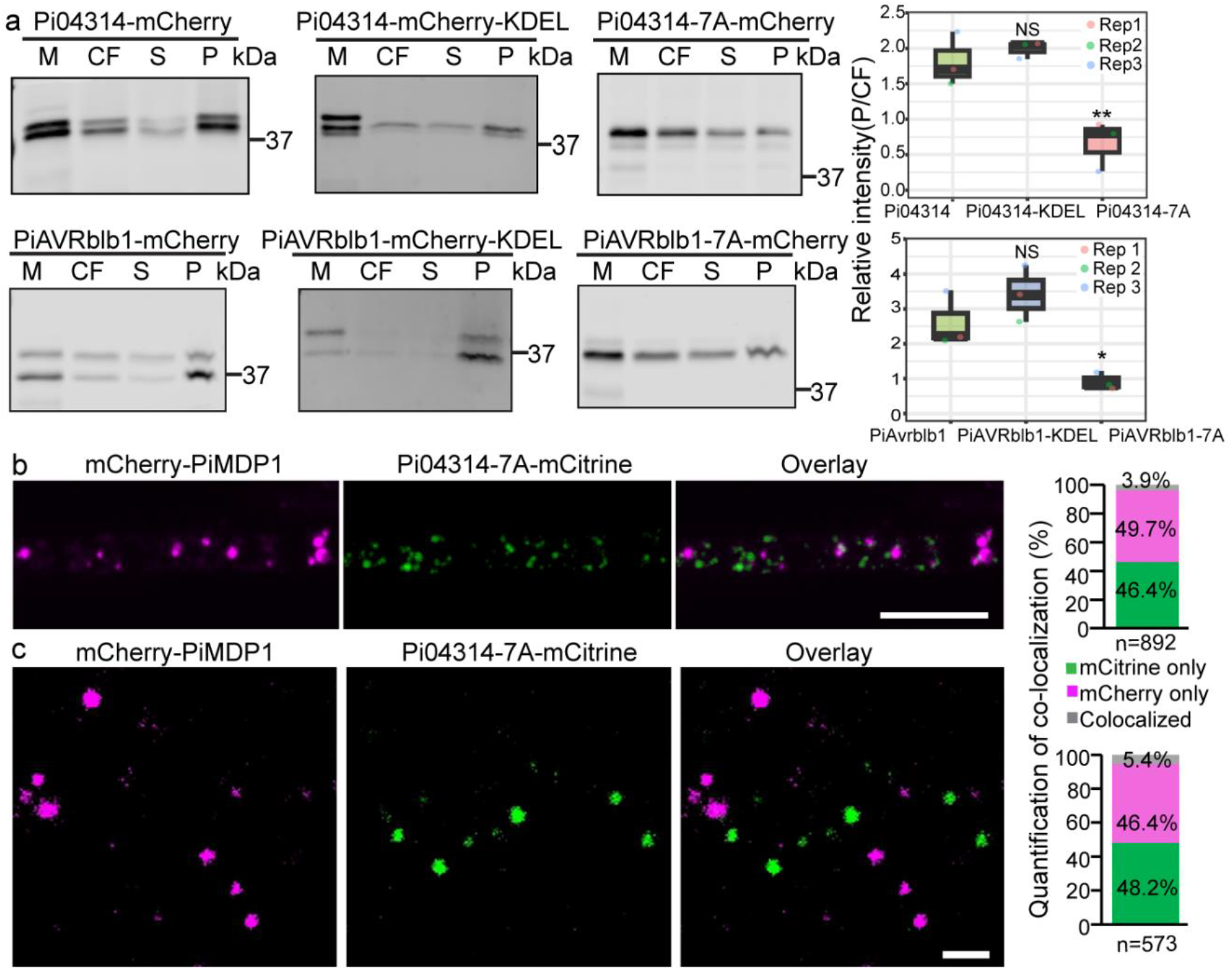
RXLR-EER motifs determine the association of effectors with extracellular vesicles (EVs) **a**, Immunoblot analysis of protein preparations of different fractions from cultured hyphae shows that RXLR effector fusions Pi04314-mCherry, PiAVRblb1-mCherry and their KDEL-modified variants are predominantly enriched in the pellet (P) fraction which contains EVs, whereas the RXLR-EER mutant variant displays markedly reduced association with the pellet fraction. Quantification of protein levels in the pellet fraction relative to that in the culture filtrate (CF) is shown in the accompanying boxplot. Data are expressed as mean ± SD from three independent biological replicates, with individual values represented by solid circles. Statistical analysis was performed using ordinary one-way ANOVA with Dunnett’s multiple comparisons test to determine significance. M= mycelia; S=supernatant. Protein sizes are indicated in kDa. **b**, Confocal microscopy images show that mCherry-PiMDP1 and Pi04314-7A-mCitrine do not colocalize in cultured hyphae. Scale bars, 10 µm. **c**, Endosomal vesicles isolated from the same dual transformants as in **b** confirm the absence of colocalization of Pi04314-7A and PiMDP1. Quantitative analysis of colocalization is presented in the accompanying graphs. The percentage of the total numbers of puncta (n) that were colocalized was measured using the DiAna plugin in Fiji (**b, c**). Scale bars, 2 µm.

### The RXLR-EER motif governs the association of RXLR effectors with EVs

Our recent work suggested that RXLR effectors such as Pi04314 and PiAVRblb1 are associated with pellets containing EVs^19^. Pi04314 was also shown to co-localize, prior to secretion, with two candidate EV markers, PiMDP1 and PiMDP2 in hyphal endo-vesicles^19^. To investigate whether the RXLR-EER motif governs this co-association, we examined WT and RXLR-EER mutant versions of the effectors (Pi04314-7A and PiAVRblb1-7A) using EVs isolated from cultured mycelia and western blotting. Although the mutants were still detected in pellets containing EVs, their abundance that relative to total secretion (CF) was significantly reduced compared to the WT proteins, indicating diminished association with EVs (Fig. 4a). In contrast, the addition of a C-terminal KDEL motif to the WT Pi04314 or PiAVRblb1did not affect EV association (Fig. 4a), consistent with our findings that KDEL does not alter the secretion route of RXLR effectors (Fig. 2a, b).

To further assess the role of the RXLR-EER motif in EV targeting, Pi04314-7A was co-expressed with the EV marker PiMDP1 in *P. infestans*. Confocal microscopy showed no co-localization between the mutant and PiMDP1 in cultured hyphae, and they were found in distinct endo-vesicle populations (Fig. 4b, c; Supplementary Fig. 5a). In contrast, WT Pi04314 strongly co-localized with PiMDP1 in the same endo-vesicle types (Supplementary Fig. 4b, c). Together, these findings demonstrate that the RXLR-EER motif and its processing are key determinants of RXLR effector sorting into an unconventional secretion pathway associated with EV production.

## Discussion

Our study provides compelling cell biological and genetic evidence that *P. infestans* deploys distinct secretion pathways for two major classes of effector proteins—apoplastic and RXLR-type cytoplasmic effectors. Whereas apoplastic effectors are secreted via the conventional ER-to-Golgi pathway, RXLR effectors utilize an unconventional secretion route that bypasses the Golgi. Critically, we demonstrate that the RXLR-EER motif is essential for determining the unconventional secretion of these cytoplasmic effectors and their association with extracellular vesicles (EVs) (Supplementary Fig 6).

Previously, sensitivity to the inhibitor BFA was used to distinguish between *P. infestans* apoplastic and cytoplasmic (RXLR) effector secretion pathways ^15, 16^. This approach was used also to demonstrate that apoplastic and cytoplasmic effectors from the fungal pathogen of rice and wheat, *Magnaporthe oryzae*, were also secreted by conventional and unconventional pathways, respectively ^17^. However, BFA treatment can have off-target effects ^18^. As such, non-disruptive cell biological and genetic approaches are required to confirm inhibitor-based studies. We found that apoplastic effectors and RXLR effectors did not co-localise at mobile endo-vesicles in *P. infestans* hyphae (Fig 1), consistent with their secretion via different pathways. Critically, whereas addition of a KDEL motif to the C-terminus of apoplastic effectors resulted in their retention in hyphae, consistent with retrograde trafficking of them from the Golgi to the ER, addition of a KDEL motif to RXLR effectors did not prevent their secretion (Fig 2; Supplementary Fig 6). These cell biological and genetic approaches, which agree with previously published BFA results ^15,16^, offer a non-disruptive means to distinguish between conventional and unconventional secretion pathways that could be adopted to verify previous findings in other fungi such as *M. oryzae* ^17^ and investigate effector secretion in other pests and pathogens of plants or animals.

Importantly, we identified the RXLR-EER motif as a key determinant of unconventional secretion of these effectors. Mutating the RXLR-EER motif not only altered the intracellular localization of RXLR effectors—leading them to co-localize in endo-vesicles with apoplastic effectors—but also shifted their secretion to a Golgi-dependent route (Fig. 3). When the RXLR-EER motif was mutated, effectors were retained in the ER upon addition of a KDEL tag, confirming they were secreted conventionally via the Golgi (Supplementary Fig. 3). We thus argue that the RXLR-EER motif acts as an ER sorting signal to an unconventional, Golgi-bypass secretion pathway. Importantly, secretion of the WT RXLR effector Pi04314-mRFP by the plant *N. benthamiana* followed a conventional ER-to-Golgi pathway that was BFA-sensitive and prevented by addition of a KDEL motif (Supplementary Fig. 3). As we showed previously that there is no efficient proteolytic cleavage of Pi04314 during secretion by *N. benthamiana*^14^ this strongly implicates such processing of the RXLR-EER motif as a requirement for sorting into an unconventional secretion pathway. Interestingly, our recent observation that RXLR effectors are often associated with extracellular vesicles (EVs), whereas apoplastic effectors are not^19^, provides further evidence that these effector classes follow distinct secretion pathways. Our data here demonstrate that the RXLR-EER motif also governs the association of RXLR effectors with EVs. Whereas WT RXLR effectors co-localized with EV markers and were enriched in EV pellets, their RXLR-EER mutants showed diminished EV-association and failed to co-localize with EV markers (Fig. 4, Supplementary Fig. 5). This suggests that the RXLR-EER motif has a critical role in the intracellular sorting of RXLR effectors toward an unconventional pathway associated with EV production (Supplementary Fig 6).

The *Phytophthora* RXLR effectors have been described as analogous to PEXEL effectors delivered into human cells by the distantly-related apicomplexan malaria pathogen *Plasmodium falciparum* ^32^. PEXEL effectors contain a related motif, RxLxE/D/Q, which is also proteolytically processed during secretion ^33^. The precise role of this motif remains unclear. Our findings reveal a role for the RXLR-EER motif in directing RXLR effectors to an unconventional secretion pathway that bypasses the Golgi following proteolytic cleavage. This is different to PEXEL effectors from *Plasmodium* which, following proteolytic cleavage in the ER, are secreted through the ER-to-Golgi pathway ^34^. Future studies should focus on identifying the molecular machinery that processes RXLR-EER motifs within *Phytophthora* and mediates their sorting into an unconventional secretion pathway associated with EV production. This machinery could represent a novel target for disease control.

## Methods

### *Phytophthora infestans* growth and plant growth

*Phytophthora infestans* (*P. infestans*) strain 3928A, along with its derived transgenic lines, was cultured on Rye Agar medium supplemented with Ampicillin (100 µg/ml) and Pimaricin (0.02%). Geneticin (10 µg/ml) was included to ensure the selection and maintenance of transgenic lines. *Nicotiana benthamiana* and the susceptible potato cultivar Solanum tuberosum ‘Maris Piper’ were grown in controlled glasshouse conditions under a 16-hour light/8-hour dark cycle.

To assess the secretion efficiency of *P. infestans*. A 14-day old plate was flooded with 3ml amended lima bean (ALB) liquid media supplemented with the appropriate antibiotics scraped with a sterile spreader and grown for 1 days in the dark without shaking. Mycelia were transferred into fresh culture tubes containing 1.5 ml ALB (with appropriate antibiotics) which had been pre-cleared by ultracentrifugation at 120 000 × g for 2 hours. Cultures were grown for a further day.

For EV/Endo-vesicle isolation. At least two 14-day old plates were flooded with 15ml amended lima bean (ALB) liquid media supplemented with the appropriate antibiotics. Sporangia were collected by pouring the suspension into a sterile culture bottle through a 70 μm cell strainer. Sporangia were allowed to grow for 2 days (or 4 days for endo-vesicle isolation) in the dark without shaking. To isolate EVs, mycelia were transferred into fresh culture bottles containing 15 ml minimal media and 15 ml ALB which had been pre-cleared by ultracentrifugation at 120, 000*g* for 2 h. Cultures were grown for a further 2 days.

### Construction of *P. infestans* transgenic lines

The vector pmCitrine:mCherry, designed for co-expression of proteins tagged with mCitrine and mCherry, was constructed by Breen et al., (2025)^19^. The coding sequence for PiSCR74 was amplified from a published vector^35^. Coding sequences for PiEPIC1 (PITG_09169) and Pi04314 (PITG_04314) were amplified from published vectors constructed by Wang et al., (2017, 2018)^15, 16^, and PiAVRblb1 (PITG_21388) and all RXLR-EER mutated versions were amplified from published vector generated by Xu et al., (2025)^14^, using primers listed in Supplementary Table 1. KDEL motif was synthesised with restriction enzyme sites (Supplementary Table 1). These coding sequences were cloned as C-terminal fusions to the fluorescent proteins, driven by the oomycete Ham34 promotor (Ham34P)^36^, using restriction enzyme cloning and standard molecular biology techniques. Plasmids were sequenced to verify the correct insertion of target sequences and prepared for downstream applications using the Promega PureYield™ Plasmid Maxiprep System. Transformation of *P. infestans* isolate 3928A was conducted following the protocol described by Welsh et al. (2025)^37^.

### Construction of expression vectors and transient expression *in planta*

The *P. infestans* genes encoding Pi04314, PiSCR74, or PiAVRblb1, including their native predicted signal peptides and C-terminally fused mCherry, with or without a KDEL motif, were amplified from above constructed vectors used in *P. infestans* transformations. The vector lacking the signal peptide (NSP-Pi04314-mRFP) was generated by Wang et al. (2017)^15^, while the NSP-PiAVRblb1-mRFP construct was produced by Xu et al. (2025)^14^. Gene-specific primers containing flanking Gateway recombination sites were used for PCR amplification (Supplementary Table 1). The purified PCR products were recombined into the entry vector pDONR201 (Invitrogen) to generate entry clones. These entry clones were subsequently recombined with the binary vector pK7WG2 and transformed into *Agrobacterium tumefaciens* strain GV3101 for transient expression in *Nicotiana benthamiana*^38^. *A. tumefaciens* cultures were infiltrated at an OD_600_ of 0.05. Brefeldin A (BFA) was applied at a final concentration of 20 µg/mL, 30–60 minutes prior to imaging, as previously described Wang et al., (2017)^15^.

### Immunoblotting and quantification

For protein extraction from *P. infestans* mycelia, approximately 0.01 g of fresh mycelium was collected in a 1.5 ml Eppendorf tube and mixed with 80 µl of 2× SDS buffer (to assess the overall secretion efficiency, all obtained mycelia were collected in a 1.5 ml Eppendorf tube and ground with a pestle in 150 µl of 2× SDS buffer.). The sample was vortexed for 1 minute and subsequently boiled at 95°C for 10 minutes. For protein extraction from culture filtrate and supernatant, total proteins were isolated using the chloroform/methanol precipitation method^39^. Briefly, 4 ml of methanol, 1 ml of chloroform, and 4 ml of sterile water were added to 1 ml of liquid samples. To assess the overall secretion efficiency, all resulted culture filtrate (∼1.5 ml) was used for precipitation. The mixture was thoroughly vortexed and centrifuged at 5000 × g for 20 minutes. The precipitated proteins were washed with 3 ml of absolute methanol, and the air-dried pellet was resuspended in 80 µl of 2× SDS loading buffer.

Samples were loaded onto 10% SDS-PAGE gels and transferred onto nitrocellulose membranes. Membranes were blocked for one hour at room temperature in 4% milk dissolved in 1× PBST (0.05% Tween 20). Primary antibodies targeting RFP/mCherry (Chromotek, 58F) and GFP/mCitrine (Roche, 11814460001) were diluted to 1:4000 and 1:2000, respectively, in blocking buffer and incubated with the membranes overnight at 4°C or room temperature for 2 hours. Following incubation, membranes were washed five times (10 minutes for each time) with 1× PBST. Secondary antibodies—IRDye® 680RD Goat anti-Rat IgG, IRDye® 800CW Goat anti-Mouse IgG (LI-COR Biosciences)—were added at a 1:5000 dilution in blocking buffer. Membranes were incubated with secondary antibodies for one hour at room temperature in the dark. Signal detection and quantification were performed using the LI-COR ODYSSEY® DLx imaging system and LICOR Image Studio™ software.

### Endo-vesicles isolation from *P. infestans*

Endo-vesicles from *P. infestans* were isolated using 1–2 g of mycelia homogenized in an equal volume of isolation buffer (150 mM Na-HEPES, pH 7.5, 10 mM EDTA, 10 mM EGTA, 17.5% [w/w] sucrose, 7.5 mM KCl, 10 mM DTT, 1× PIC-W, 1× PIC-D, and 1× E-64) as described by Wang et al. (2023)^40^. For samples designated for Triton X-100 treatment, the supernatant was divided into two equal volumes. One portion was treated with Triton X-100 to a final concentration of 1%, while the other received an equivalent volume of sterile distilled water as a control. All samples were thoroughly mixed and incubated on ice for one hour. After incubation, the samples were adjusted to 40.6% (w/w) sucrose by mixing with 62% (w/w) sucrose solution. A 2.4 mL aliquot of this sucrose mixture was loaded at the bottom of a 13.2 mL ultracentrifuge tube and overlaid sequentially with 3 mL of 35% (w/w) sucrose, 3 mL of 25% (w/w) sucrose, and 3 mL of 8.5% (w/w) sucrose to form a gradient. The loaded gradient was centrifuged at 400,000*g* for 3 hours at 4 °C using a Beckman Coulter SW 41 Ti rotor. The two interfaces between the 8.5% and 25%, 25% and 35% sucrose layers was carefully collected and pelleted by ultracentrifugation at 100,000*g* for 1 hour at 4 °C. Pellets were prepared for microscopy analysis and immunoblotting. All sucrose solutions were prepared in 150 mM Na-HEPES buffer (pH 7.5) and included protease inhibitors (1× PIC-W, 1× PIC-D, and 1× E-64) to ensure vesicle integrity.

### Transmission microscopy (TEM)

25 μl of endo-vesicle samples or buffer control was mixed with 25 μl of 4% paraformaldehyde (PFA) in 0.1 M sodium phosphate pH 7.4 to give a final concentration of 2% PFA. 30 μl aliquots were pipetted onto Parafilm with nickel grids (carbon-coated, formvar, 300 mesh) placed on top, and incubated at room temperature for 30 minutes. Grids were washed with 1x PBS twice and incubated for 10 minutes with 50μl 1% glutaraldehyde. Grids were then washed with 100 μl sterile water seven times and stained with 4% uranyl oxalate for 10 minutes at room temperature. Grids were transferred onto two 50 μl drops of uranyl acetate (0.15% methylcellulose and 0.4% uranyl acetate) sequentially and left on the second drop for 10 minutes on ice. Excess liquid was wicked off with filter paper and grids were inspected with a JEOL 1400 at 80 kV.

### Confocal microscopy and data quantification

*P. infestans* hyphae grown in ultracentrifugation cleaned ALB medium on microscope slides were imaged at 1–2 days post-inoculation, while leaf infections on *Solanum tuberosum* were imaged at 2–3 days post-inoculation. *N. benthamiana* leaves were imaged at 2 days after *Agrobacterium* infiltration. Imaging was performed using a Nikon A1R confocal microscope.

For GFP fluorescence, the 488 nm line from an argon laser was used for excitation, and emissions were collected between 500–530 nm. For mCitrine fluorescence, the 514 nm line from an argon laser was used for excitation, and emissions were collected between 520–550 nm. For mCherry fluorescence, a 561 nm laser was used for excitation, with emissions collected between 600–630 nm. The pinhole was set to 1.2 airy unit to optimize resolution. Images were processed using OMERO, or ImageJ software. To assess the effector secretion efficiency during infection, haustoria and adjacent area of invasive hyphae were chosen from single optical sections for measurement. The Fiji plugin DiAna was used for endo-vesicle co-localization analyses^41^.

### EV isolation from *P. infestans* culture

EVs were isolated from *P. infestans* culture as described before^19^. Approximately 30 ml culture was used for EV isolation. 1 ml of culture filtrate (CF) was collected after the removal of mycelia using a 70 μm cell strainer. The filtered CF was centrifuged for 10 minutes at 2000*g* and filtered through a 0.45 μm syringe filter, following a further spin at 10,000*g* for 30 minutes to remove all cellular debris and large vesicles. Supernatant from previous step was then ultracentrifuged at 100,000*g* for 1 hour at 4 °C (Beckman Coulter Type50.2 Ti rotor) to finally obtain the crude EV pellet (P) samples, pellet was dissolved in 80 μl SDS buffer for western blot.

## Acknowledgements

We are grateful to colleagues Drs Hazel McLellan and Susan Breen for helpful discussions during the development of this research. We thank Dr Kasia Mackinnon for technical support. The research was supported by United Kingdom Research and Innovation Horizon Europe Guarantee Marie Skłodowska-Curie Actions Postdoctoral Fellowship (EP/Z00196X/1); Biotechnology and Biological Sciences Research Council grants BB/Z515115/1 and BB/Z517495/1; European Research Council (ERC)-Advanced grant PathEVome (787764); and Scottish Government Rural and Environment Science and Analytical Services Division (RESAS) JHI B1-1.

## Author contributions

S.W., W.W and P.R.J.B. conceived of and designed the research; S.W. and W.W. performed the research; L.X provided resources of *P. infestans* transformants; L.W. provided technical support; S.W., W.W and P.R.J.B. analysed data; S.W., P.R.J.B., S.C.W. and P.C.B. won funding for the research; and S.W., W.W and P.R.J.B. wrote the paper with input from all co-authors.

The authors declare no competing interest.

