## Supplementary Figures for "The RXLR-EER Motif Determines an Unconventional Secretion Pathway Associated with Extracellular Vesicle Production"

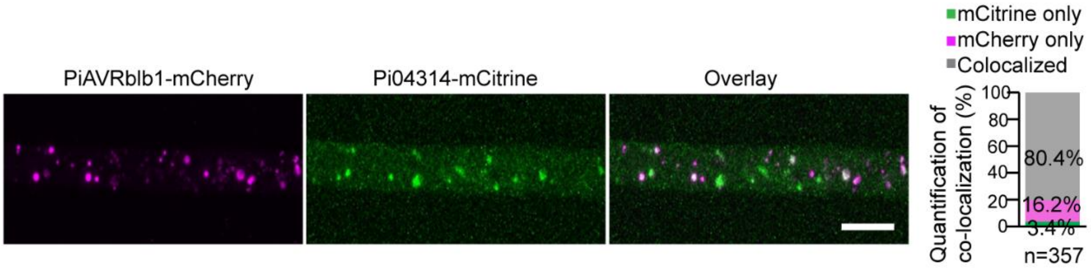


**Supplementary Fig. 1| Two RXLR effectors are colocalized in cultured hyphae.** Confocal microscopy images show that PiAVRblb1-mCherry and Pi04314-mCitrine are colocalized in cultured hyphae. Scale bars, 10 µm. Quantitative analysis of colocalization is presented in the accompanying graph. The percentage of the number of total (n) puncta that were colocalized was determined using the DiAna plugin in Fiji.

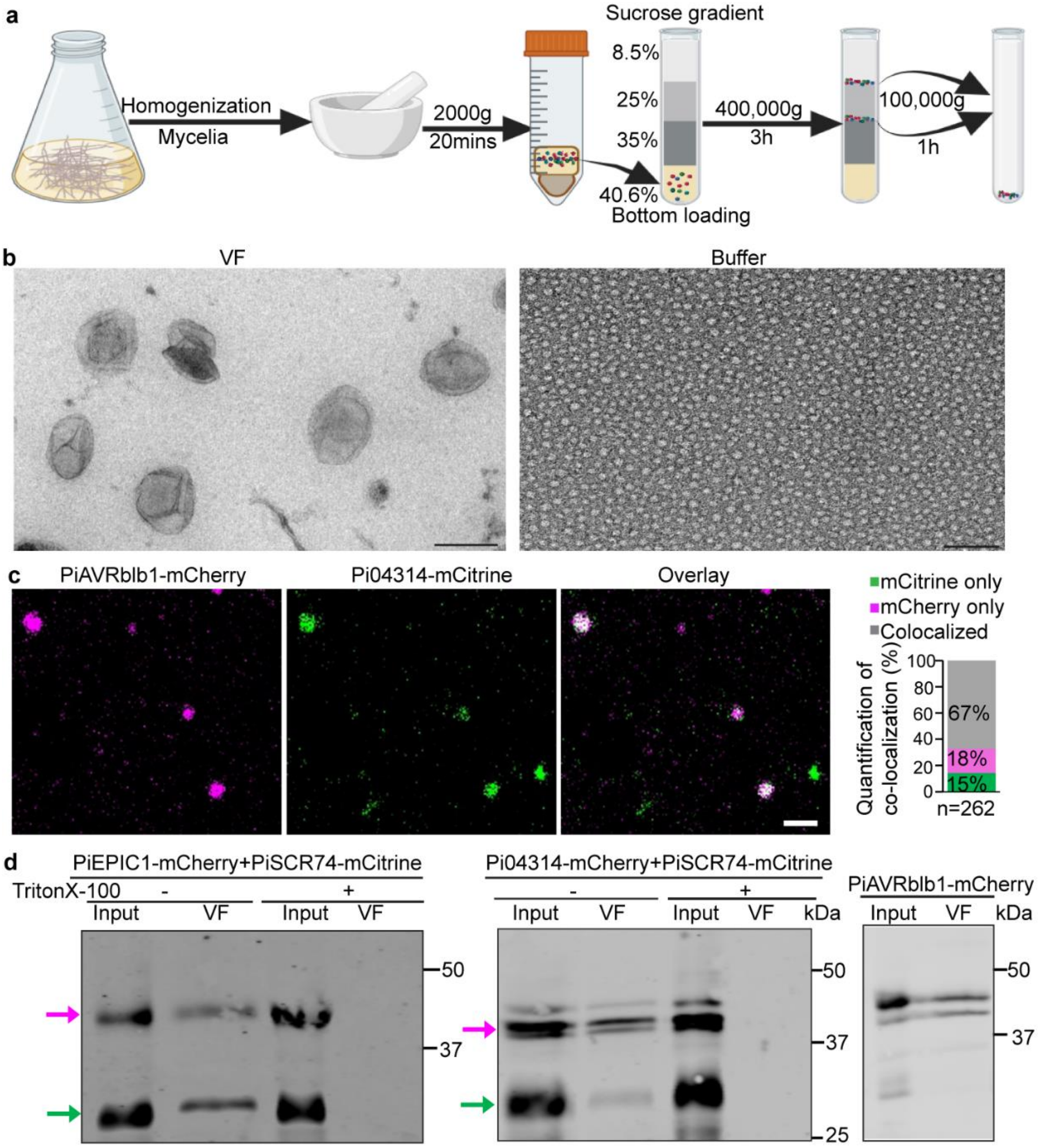


**Supplementary Fig. 2| Two RXLR effectors are colocalized in isolated endo-vesicles isolation. a**, Schematic representation of the process used to isolate endo-vesicles from *P. infestans* transformants. **b**, Transmission electron microscopy reveals that the vesicle fraction (VF) contains characteristic membrane-bound vesicles. The buffer used for endo-vesicle isolation served as a negative control and contained no such structures. Scale bar, 200 nm. **c**, Endosomal vesicles isolated from the same dual transformants in **Supplementary Fig 1** confirm the colocalization between two RXLR effectors. Quantitative analysis of colocalization is presented in the accompanying graph. The percentage of the number of total (n) puncta that were colocalized was determined using the DiAna plugin in Fiji. Scale bars, 2 µm. **e**, Immunoblot analysis shows that both apoplastic and RXLR effectors are detected in vesicle fractions. "Input" refers to the supernatant collected after centrifugation at 2,000 × *g*. (+) indicates samples treated with Triton X-100 detergent, while (–) indicates untreated controls. Magenta arrows denote mCherry-tagged proteins; green arrows indicate mCitrine-tagged proteins. Protein sizes are indicated in kDa.

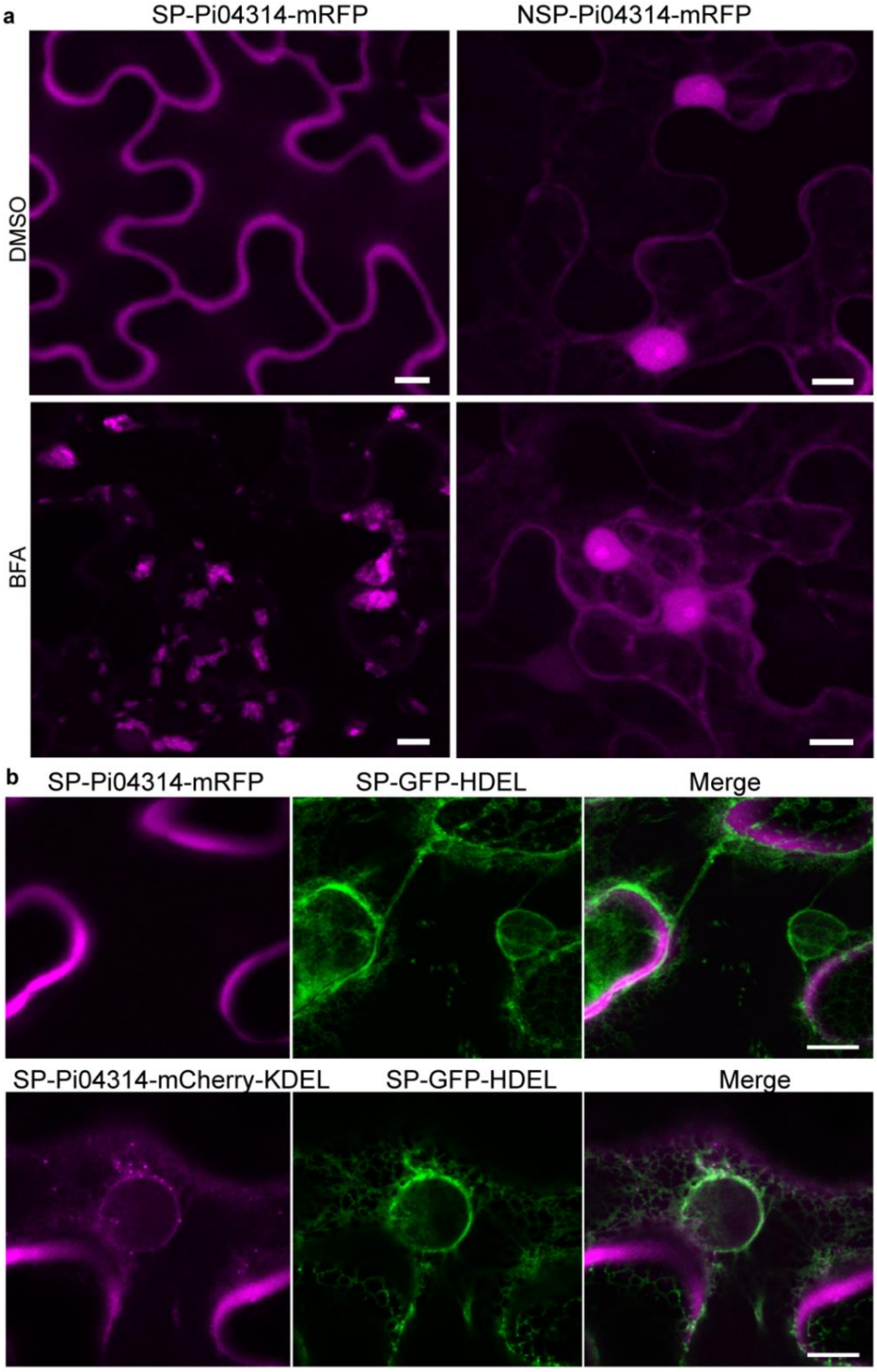


**Supplementary Fig. 3| Cleavage of RXLR-EER motif is required for unconventional secretion pathways. a**, Confocal microscopy images of transiently expressed effector fusion constructs show that the secretion of the RXLR effector Pi04314 with a signal peptide (SP) is sensitive to Brefeldin A (BFA) treatment in *Nicotiana benthamiana*, while DMSO-treated samples serve as a negative control. Pi04314 -mRFP expressed without the signal peptide (NSP) localizes predominantly to the nucleus and nucleolus, and this localization is unaffected by BFA treatment. Scale bars, 10 µm. **b**, Confocal microscopy images show that the fusion protein SP-Pi04314-mCherry-KDEL is retained in the plant endoplasmic reticulum (ER), in contrast to the SP-Pi04314-mRFP, which is secreted. Transgenic *N. benthamiana* expressing SP-GFP-HDEL was used to confirm ER localization. Scale bars, 10 µm.

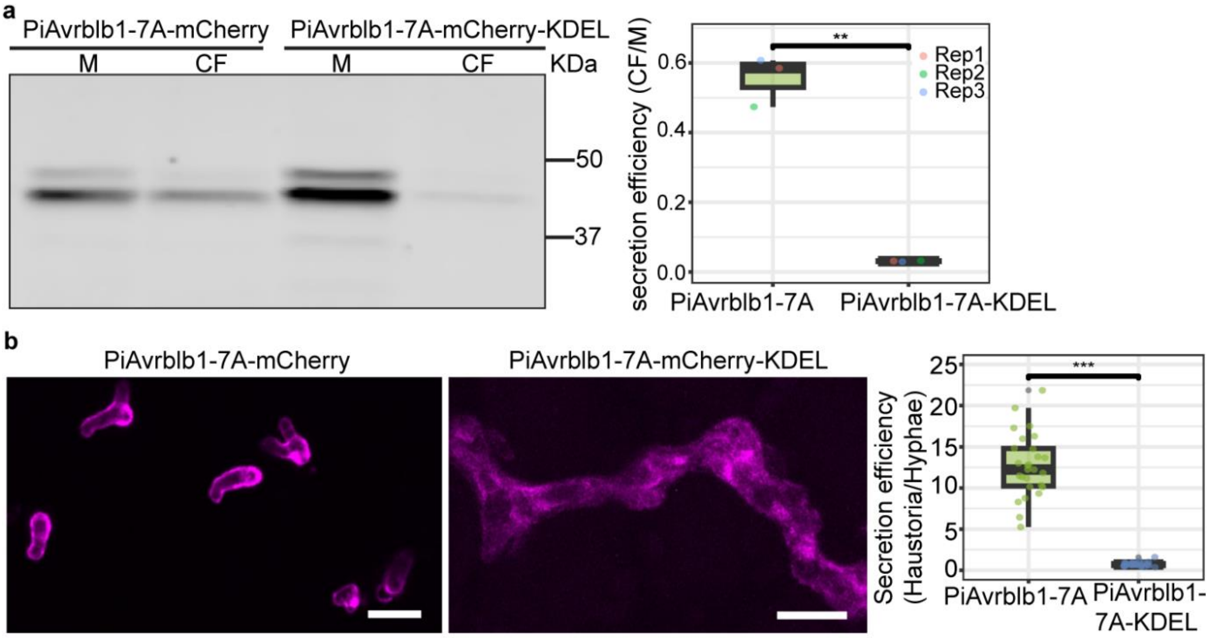


**Supplementary Fig. 4| ER retention of KDEL-tagged RXLR mutant confirms secretion inhibition. a**, Immunoblot analysis shows substantial intracellular accumulation of PiAVRblb1-7A-mCherry-KDEL in the mycelia (M), contrasting with the level of secretion observed for PiAVRblb1-7A-mCherry. Secretion efficiency was quantified by calculating the ratio of protein abundance in the culture filtrate (CF) relative to that in M. The corresponding boxplot presents the quantification, with data shown as mean ± SD from three biological replicates, each indicated by individual solid circles. Protein molecular weights are labeled in kDa. **b**, Confocal microscopy of infectious hyphae in potato leaf tissue reveals markedly reduced haustorial accumulation of PiAVRblb1-7A-mCherry *in planta* when it is fused to the KDEL motif, compared to the unmodified construct. Quantification of fluorescence intensity ratios between haustoria and hyphae is provided in the accompanying boxplots. Data represent mean ± SD from three independent biological replicates, with individual values shown as solid dots. Statistical analysis was performed using T-test. Scale bars, 10 μm.

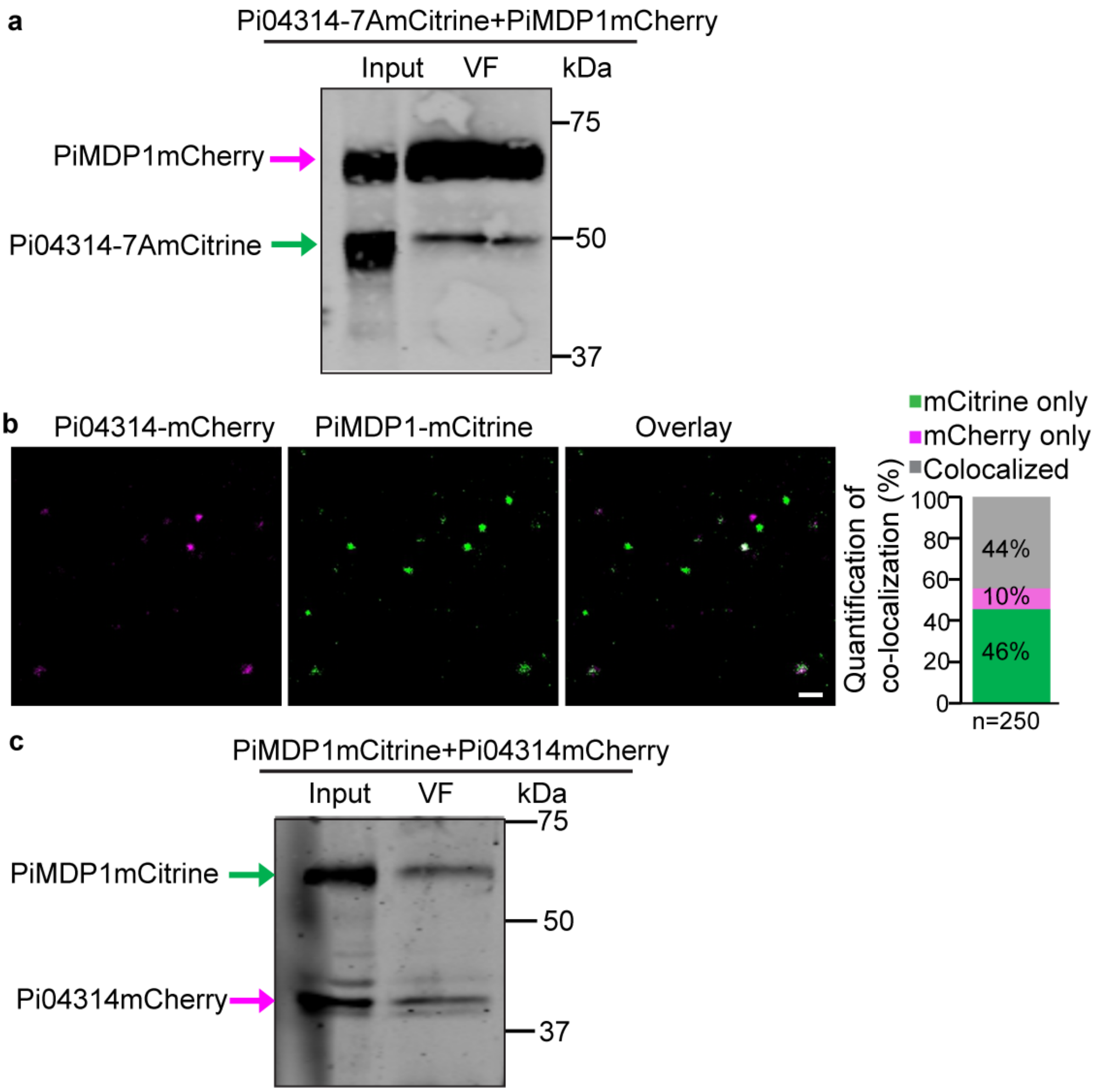


**Supplementary Fig. 5| The EV marker PiMDP1 colocalizes with the RXLR effector, but not with its mutated variant. a**, Immunoblot analysis shows that both RXLR mutated effectors and PiMDP1 fusions are detected in endo-vesicle fractions (VF). **b**, Endosomal vesicles isolated from the dual *P. infestans* transformant expressing Pi04314-mCherry and PiMDP1-mCitrine confirm colocalization between the RXLR effector and the EV marker PiMDP1. Quantitative analysis of colocalization is presented in the accompanying graph. The percentage of the number of total (n) puncta that were colocalized was determined using the DiAna plugin in Fiji. Scale bars, 2 µm. **c**, Immunoblot analysis shows that both the wild-type RXLR effector and PiMDP1 fusions are present in the vesicle fractions. “Input” refers to the supernatant collected after centrifugation at 2,000 × *g* (**a**, **c**). Protein sizes are indicated in kDa (**a**, **c**).

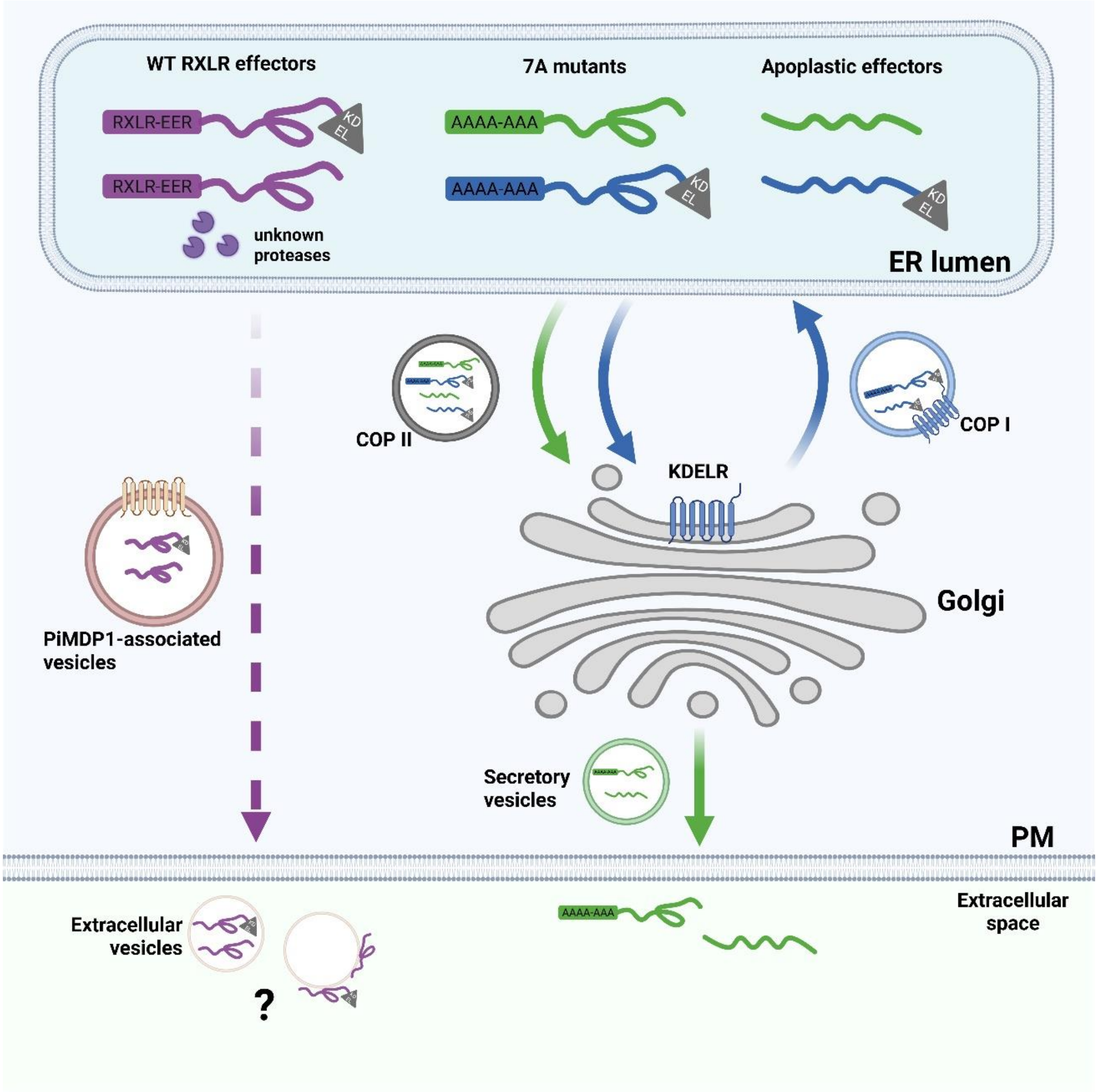


**Supplementary Fig. 6| Summary Diagram of the findings in this work.** Cytoplasmic effectors containing the RXLR-EER motif (purple) are proteolytically processed [14] prior to secretion via an unconventional pathway (dashed purple line), in association with the transmembrane marker PiMDP1 [19], that bypasses the Golgi. Addition of a KDEL motif does not prevent secretion of WT RXLR effectors in association with uncharacterised (question mark) extracellular vesicles [19]. In contrast, mutation of the RXLR-EER motif to alanines (7A mutant) results in co-secretion of RXLR effectors with apoplastic effectors (green arrows) via Golgi. Addition of a KDEL motif to either the 7A mutant RXLR effector or to an apoplastic effector (blue arrows), results in their retention in the ER due to KDEL receptor (KDELR)-mediated retrograde trafficking from the Golgi. PM is plasma membrane.
